# Change point detection with multiple alternatives reveals parallel evaluation of the same stream of evidence along distinct timescales

**DOI:** 10.1101/2020.11.29.403139

**Authors:** Alexa Booras, Tanner Stevenson, Connor N. McCormack, Marie E. Rhoads, Timothy D. Hanks

**Affiliations:** Center for Neuroscience, University of California Davis, Davis, CA, USA; Department of Neuroscience, University of California Los Angeles, Los Angeles, CA, USA; Department of Neurology, University of California Davis, Sacramento, CA, USA

## Abstract

In order to behave appropriately in a rapidly changing world, individuals must be able to detect when changes occur in that environment. However, at any given moment, there are a multitude of potential changes of behavioral significance that could occur. Here we investigate how knowledge about the space of possible changes affects human change point detection. We used a stochastic auditory change point detection task that allowed model-free and model-based characterization of the decision process people employ. We found that subjects can simultaneously apply distinct timescales of evidence evaluation to the same stream of evidence when there are multiple types of changes possible. Informative cues that specified the nature of the change led to improved accuracy for change point detection through mechanisms involving both the timescales of evidence evaluation and adjustments of decision bounds. These results establish three important capacities of information processing for decision making that any proposed neural mechanism of evidence evaluation must be able to support: the ability to simultaneously employ multiple timescales of evidence evaluation, the ability to rapidly adjust those timescales, and the ability to modify the amount of information required to make a decision in the context of flexible timescales.

## Introduction

Decision making often involves evaluation of information from the environment to form a response. In order to behave appropriately in a rapidly changing world, individuals must be able to detect when changes occur in that environment. This process is referred to as change point detection^1^. In real world situations, there are many possible aspects of the environment that could change, but there are often constraints on what changes are possible. For instance, in tracking a fly in the air, the fly could change direction along three dimensions, but in tracking an ant on the ground, the ant could change direction along only two dimensions. We sought to address how knowledge about the space of possible changes affects human change point detection.

Sequential sampling models that have been applied to a wide range of decision making tasks provide a framework to describe the process of change point detection. In these models, an observer makes a choice based on a decision variable that combines information from different samples of evidence gathered over time, a process we refer to as evidence evaluation. Evidence evaluation can take a variety of forms, and the optimal form depends on task demands and environmental conditions. For example, in a stable environment, all samples of evidence can be treated equally, and evidence can be perfectly integrated over a long timescale to maximize performance^2–7^. In contrast, in an unstable environment, samples from the past are less likely to reflect the current state of the world, and evidence should be evaluated over a briefer timescale to avoid integrating information that is no longer relevant^8–10^. Change point detection is a specific example that involves an unstable environment and shorter timescales of evidence evaluation^10–12^.

Knowledge about the space and type of possible changes may affect evidence evaluation for change point detection. Different types of changes may have distinct statistical properties that demand different optimal timescales of evidence evaluation. For example, detecting the approach of a speedy predator benefits from a shorter timescale of evidence evaluation than detecting the approach of a slow and stealthy predator. Thus, information about the nature of the change -- for example being in a location with only speedy predators or only slow, stealthy predators -- has the potential to improve change detection performance through optimization of the timescale of evidence evaluation.

Knowledge about the space and type of possible changes may also improve change detection performance through mechanisms typically associated with selective attention^13–15^. In the sequential sampling framework, each sample of evidence that is evaluated carries a signal about the environment that is potentially corrupted by signal-independent variability known as noise^2,3^. That noise may involve environmental factors, but it may also be related to neural processing^16–19^. To the extent the brain accounts for this noise, knowledge about the possible changes may affect the signal to noise ratio of the evidence evaluated for change point detection.

Numerous studies have described improvements in performance with attention in a variety of tasks that are attributable to increases in the signal to noise ratio that enhance perceptual sensitivity of attended stimuli^20–22^. Thus, knowledge about the space of possible changes may alter evidence evaluation by changing perceptual sensitivity for change point detection.

A third possibility is that knowledge about the space and type of possible changes may alter decision commitment policies. In the sequential sampling framework, a bound on the decision variable determines the timing of decision commitment^23–26^. For change point detection, a lower decision bound results in fewer misses at the expense of more false alarms, and a higher decision bound results in fewer false alarms at the expense of more misses^11^. Thus, the decision bound determines the tradeoff between false alarms and misses. There are many important considerations for setting the decision bound for change point detection. Among those is the number of potential changes that are simultaneously being monitored in any given situation. If each potential change carries a fixed probability of false detection, then the greater the number of potential changes simultaneously under consideration, the greater the probability of a false alarm detection of one of those potential changes. Therefore, higher decision bounds may be required in situations with a greater number of potential changes that need to be considered. Following this logic, if fewer potential changes are under consideration, false alarm rates can be kept in check even with relatively lower decision bounds with the benefit of also reducing the miss rate. Thus, knowledge about the space of possible changes may affect the setting of decision bounds for change point detection.

Previous research suggests that both perceptual and decision components of processing can be influenced by cues about possible changes in the environment^27^. As a particularly relevant example for our study, Sridharan and colleagues showed dissociations of the effects of spatial cueing between sensory and decision components of processing for “yes/no” change detection in the visual domain, where people have to report if a change has occurred between two comparison stimuli. In particular, they found that sensory effects of cuing were spatially localized while decision effects were not^28^. In order to distinguish these possibilities, they used a multidimensional signal detection model that relies on a comparison of two samples of evidence^29^, as is appropriate for the “yes/no” detection task they studied. However, it is not known how cues about the space and type of possible changes influences change point detection, which involves evaluation of multiple sequential samples of evidence rather than a comparison of two.

Here, we sought to address how knowledge about the space of possible changes affects evidence evaluation and decision bounds for an auditory change detection task we have developed. In this task, subjects must rapidly report when a change occurs in an auditory stimulus that is unpredictable in time. The change in the stimulus could be of two different varieties, and we provided cues that either informed the subject about the type of change that would occur or were uninformative about the type of change. We compared performance when the cue was informative versus uninformative using model-free psychophysical reverse correlation analyses combined with sequential sampling models to characterize behavior. We found that subjects can simultaneously apply distinct timescales of evidence evaluation to the same stream of evidence. Furthermore, accuracy for detection improved with informative cues through mechanisms involving these timescales of evidence evaluation and adjustments of decision bounds. These results establish important capacities of information processing for decision making that any proposed neural mechanism of evidence evaluation must be able to support.

## Methods

### Subjects

We recruited 15 subjects to perform this experiment. Six subjects elected to discontinue the experiment and we analyzed the data of the remaining nine subjects. All subjects were undergraduate students from UC Davis over the age of 18. None had any prior knowledge of the research motivations or task design prior to data collection. Subjects were compensated with $10 Amazon gift cards for every one hour session, regardless of their performance on the task. The study procedures were approved by the UC Davis Institutional Review Board and all subjects provided informed consent.

Subjects performed one session of the experiment per day, with each session consisting of three blocks lasting 15 minutes and separated by short breaks. The first session was used as a training session and the data was not analyzed. Subjects completed either three or four non-training sessions, with three subjects electing not to return for the final session.

### Change Detection Task

Subjects were asked to perform an auditory change detection task. They were seated in an isolated room and given headphones to listen to the stimulus. The testing apparatus consisted of three ports with LEDs that indicated whether a port could be used, and IR sensors that detected when a finger was inserted into a port. The stimuli were generated with Psychtoolbox and programmed in MATLAB. The task was controlled through BPod which recorded the behavioral responses in real time. Auditory feedback was generated by the PC and played through the headphones.

Subjects were instructed to report a change in the underlying rate of a series of clicks randomly generated by a Poisson process. The change in rate could be either an increase or a decrease. The underlying rate was initially 60 Hz. For all trials the frequency changed with equal probability by ± 10, ± 30, or ± 50 Hz. There was always a minimum of 1 s before any change, after which the change time was drawn from an exponential distribution with a mean of 3.5 s and truncated at a maximum of 7.5 s. The exponential distribution results in a flat hazard rate of the stimulus change between the 1-s pre-change period and the 7.5-s maximum, so the probability of the change occurring did not vary throughout most of the trial. Before each trial, a cue on the computer screen displayed a word indicating what type of trial would occur. The cues were either informative, stating “increase” or “decrease”, or uninformative, stating “either”. The trial type was chosen at random, with a 50% chance of the trial being informed. For uninformed trials, the change direction was chosen at random with a 50% chance of the trial being an increase. This resulted in an equal amount of informed and uninformed trials, and increase and decrease trials (**Table 1**).

**Figure 1.**
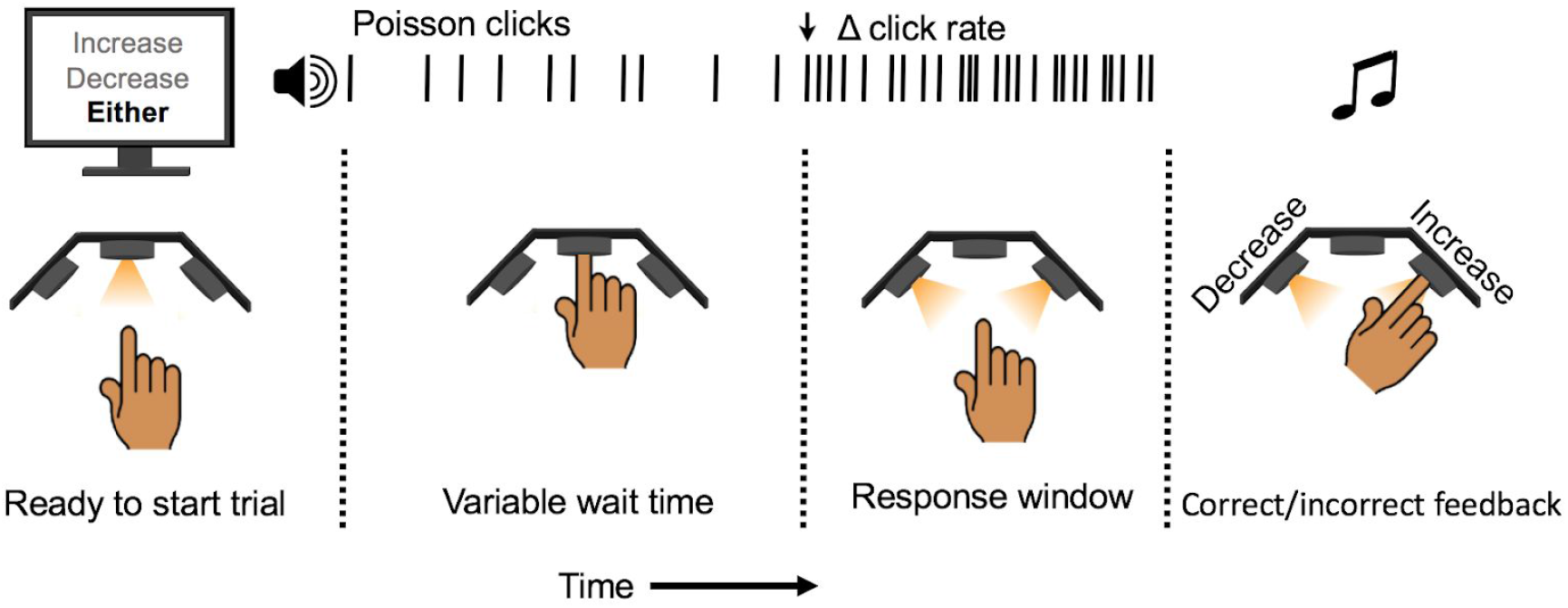
Schematic of change detection task. An illuminated LED indicates the trial is ready to begin. The computer screen displays a word indicating what type of trial will occur. The cues are either informative, stating “increase” or “decrease”, or uninformative, stating “either”. Subjects insert their finger into the port, triggering the onset of a stochastic auditory stimulus and keep their finger in the port until they detect the change in click rate upon which they remove their finger. Then they insert their finger into either the left port to report a decrease or the right port to report an increase.

**Table 1.** Trial condition countsfor each subject and totalsfor combined subject analysis.

| Subject | Informed Increase | Informed Decrease | Uninformed Increase | Uninformed Decrease |
| --- | --- | --- | --- | --- |
| 1 | 379 | 374 | 405 | 390 |
| 2 | 555 | 559 | 556 | 532 |
| 3 | 451 | 428 | 475 | 463 |
| 4 | 541 | 561 | 544 | 600 |
| 5 | 510 | 526 | 511 | 533 |
| 6 | 551 | 545 | 511 | 510 |
| 7 | 390 | 400 | 419 | 427 |
| 8 | 345 | 364 | 372 | 380 |
| 9 | 415 | 410 | 440 | 417 |
| **Combined** | **4137** | **4167** | **4233** | **4252** |

At the start of each trial, an illuminated LED in the center port of the apparatus indicated the trial was ready to begin and the trial cue appeared on screen. Subjects inserted their finger into the port to start the trial, triggering the onset of the auditory stimulus. They were instructed to keep their finger in the port until they detected the change in click rate, at which point they were instructed to remove their finger immediately which ended the stimulus. If subjects did not remove their finger within a 0.8 s response window after the change, the stimulus would end automatically. If the subject removed their finger within the response window, side LEDs illuminated indicating that the subject should report whether they thought the change was an increase or decrease in click rate. If the subject responded to the change within the response window and correctly reported the direction of change, this trial was a “hit”. Trials in which subjects responded within the response window but reported the incorrect direction were few in quantity and were therefore not analyzed. There were two primary types of errors: premature responses, or “false alarms”, and failures to respond, or “misses”. In the case of false alarms, subjects were instructed to report the direction that they actually responded to, even if this was different than what the cue indicated. In the case of misses, the side LED corresponding to the correct direction of change briefly illuminated at the end of the trial. Immediately following the direction report, feedback was given via a high or low pitched auditory tone to indicate success or failure on that trial respectively. There were two additional types of errors that signified lapses in attention to the task--responses opposite the informative cue or responses within 0.75 s of the trial start--that were excluded from analysis.

### Psychometric Functions

Psychometric functions were computed for individual subjects and for pooled subject data. Hit rates, false alarm rates, and reaction times were calculated for each of the four trial categories: informed increase, informed decrease, uninformed increase, and uninformed decrease. Hit rates were calculated for each change delta in each category as the ratio of ‘hits’ to the number of trials where a change occurred; therefore false alarms were excluded from the calculation. 95% confidence intervals were calculated using the MATLAB *binofit* function. To quantify performance changes between uninformed and informed conditions the change delta associated with a 50% hit rate was estimated by fitting a sigmoid curve to the hit rates at each change delta in each trial category. Confidence intervals were constructed for this threshold change delta with a bootstrapping method wherein thresholds were computed from hit rate curves constructed from data matching the trial count resampled 1000 times with replacement. The 95% confidence intervals were estimated as the 5th and 95th percentiles of the bootstrapped thresholds.

Informed false alarm rates were calculated as the ratio of informed false alarms to the total number of trials in each informed condition. False alarms with responses opposite the cue were excluded from the calculation. Uninformed false alarm rates in each direction were calculated as the ratio of false alarms in one direction to the total number of uninformed trials minus the false alarms in the other direction. This exclusion attempts to account for the possibility that a false alarm in one direction masks the potential for a later false alarm in the other direction. 95% confidence intervals were calculated using the MATLAB *binofit* function. Confidence intervals for false alarm rate differences between conditions were calculated using a bootstrapping method where rate differences were calculated from data matching the trial count resampled 1000 times with replacement. The 95% confidence intervals were estimated as the 5th and 95th percentiles of the bootstrapped differences.

Reaction times were calculated as the time from the click rate change to the subject’s response. The 95% confidence intervals were estimated as the 5th and 95th percentiles of 1000 bootstrapped means.

### Psychophysical Reverse Correlations

We utilized psychophysical reverse correlations (RC) of false alarms to investigate how subjects evaluate evidence in different trial conditions^11,30–32^. RC “kernels” were produced by aligning false alarm trial stimuli to the time of the response, computing instantaneous click rates by convolving the click times with a half gaussian filter (σ = 0.075 s), and averaging over all trials grouped by cue and response direction. False alarms with responses opposite the cue were excluded from the calculation. To quantitatively compare the RC kernels, we computed the average change in click frequencies from baseline for each category of false alarms. This was done by counting the number of clicks in the 1 second window preceding the false alarm and subtracting the baseline rate of 60 Hz. The 95% confidence intervals were estimated as the 5th and 95th percentiles of 1000 bootstrapped values. We also estimated the timescale of evidence evaluation by computing the start of the RC kernel in each trial type. Kernel starts and confidence intervals were estimated by fitting a piecewise linear function from the baseline to the kernel start and from the kernel start to the kernel maximum (or minimum).

### Statistical Significance

Hypothesis testing was performed using a bootstrap method to avoid assumptions about the underlying distributions^33^. Test statistics were computed from 1000 bootstrap data sets sampled with replacement from a pseudo null distribution that contain the same number of trials as the original data set. For 50% hit rate threshold and false alarm rate differences between informed and uninformed conditions, a one-sided p-value was calculated from the number of bootstrapped differences that were opposite in sign to the average differences for increasing and decreasing stimuli. For all other metrics, t-statistics were calculated based on the null hypothesis that the metrics came from the same underlying distribution, so bootstrap data sets were sampled from a pseudo null distribution containing all values from both data sets. Two-sided p-values were computed from these statistics from the number of bootstrap statistics whose absolute values were greater than the absolute value of the test statistic computed from the unmodified data set.

### Model Based Analysis

To generate a better algorithmic understanding of the differences in decision making processes between trial conditions, we fit sequential sampling behavioral models separately to informed increase, informed decrease, and uniformed trials from individual and combined subject data. In general, the model estimates a response time for each trial by convolving the clicks with an exponential filter to produce a decision variable (DV), determining when the DV crosses a bound, and adding a non-decision time (NDT) delay to account for sensory and motor delays (**Figure 5**). To account for behavioral variability, we added noise to this process and computed response time (RT) distributions. We fit our model to the behavioral data by maximizing the likelihood that the subjects’ response times could be produced by our model parameters: exponential filter width (*τ*), bounds, mean NDT (*μ_NDT_*), NDT variability (*σ_NDT_*), and process noise (*σ_process_*). Models for the informed conditions consisted of a single process with a single high or low bound, whereas the uninformed model consisted of two processes: one to detect increases and the other to detect decreases. We also compared two different versions of the uninformed model: one with separate filter widths per process and another with a shared filter width. Models for all three conditions were fit simultaneously in order to share the same NDT variability, which slightly reduced the overall model complexity.

The model uses the click stimulus to drive the dynamics of a decision variable that is governed by an Ornstein-Uhlenbeck (OU) process with input^34^:

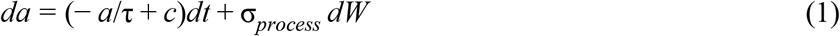

where *a* is the DV (in Hz), *c* is the DV input within each interval, *σ_process_* represents noise at the level of the DV, and *dW* is white noise. For our purposes, *c* was calculated as:

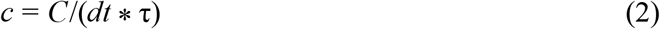

where *C* is the number of clicks in each time interval. Dividing the click frequency in the time interval (*C/dt*) by the filter width scales the DV units to Hz, which allows for direct parameter comparison across trial conditions with different filter widths.

To calculate RT distributions for each trial, we used a discretized solution to the Fokker-Planck equation corresponding to equation (1) with a timestep (*dt*) of 0.02 s. With this solution, we propagated a discretized probability distribution of DVs for each trial starting from DV = 0 at t = 0. This propagation was performed in two segments: first without bounds for 0.8 seconds, then with upper and/or lower ‘sticky’ bounds depending on the trial type. Informed increase trials used an upper bound, informed decrease trials used a lower bound, and uninformed trials used both an upper and lower bound. Sticky bounds were implemented so that any portion of the DV distribution that crossed these bounds would persist there, and thus, the proportion of the DV distribution that had crossed the bound would continually increase over time. The first DV propagation segment is designed to correspond to the period of time right after stimulus onset where subjects know there is no change and do not need to respond. This is especially important when implementing lower sticky bounds to establish a baseline DV estimate that is higher than the lower bound. We used the amount of the DV distribution that crossed the bound within each time bin as the bound crossing probability distribution, and then computed RT distributions by convolving the bound crossing distribution with a gaussian NDT filter parameterized by *μ_NDT_* and *σ_NT_*. Uninformed trials had separate RT distributions for responding to increases and decreases that were computed as joint distributions of responding to changes in one direction over the other.

The likelihood of each trial was calculated from the RT distribution by summing the distribution values in a 0.2 second window around the subject’s response time. Uninformed trials used the RT distribution corresponding to the reported direction of the response. For ‘miss’ trials where there is no response time, the trial likelihood was calculated as the remainder of the DV distribution that did not cross any bounds before the end of the trial. Optimal parameters were found by maximizing the cumulative log likelihood over all trials using both Bayesian Adaptive Direct Search (BADS)^35^ and Variational Bayesian Monte Carlo (VBMC)^36,37^ optimization algorithms. These algorithms were chosen because they were both designed to solve difficult optimization problems with non-analytical likelihood calculations, and VBMC allows for confidence interval estimation. BADS was first used to find a region of the parameter space with a high likelihood. The returned parameters were then used as the starting point for VBMC, which simultaneously estimates posterior distributions of the model parameters and a lower bound of the log model evidence (ELBO). VBMC requires a prior over the parameters, and since we made no assumptions about the parameter distributions, we used a uniform prior over the range of possible parameter values. Optimal values were estimated as the mean of 100,000 samples randomly drawn from each approximated posterior distribution. 95% confidence intervals were estimated as the 5th and 95th quantiles of the same random samples. Hypothesis testing between analogous parameters across trial conditions was performed by a two-tailed t-test using samples drawn from the fitted posterior distributions. ELBO values are analogous to the Bayes factor and were used for model selection. Multiple model fits were run with varying starting parameter values to ensure the optimal parameters were consistently found.

## Results

### Task Structure

In this experiment, nine subjects performed an auditory change detection task (see methods for details). They listened to a stream of auditory clicks generated by a 60 Hz Poisson process while holding their index finger in a response port. Subjects were told to remove their finger when they detected either an increase or decrease in the generative click rate. The timing and magnitude of the change were randomly chosen. At the start of each trial a cue either informed subjects of the direction of the change by stating “increase” or “decrease”, or the cue was uninformative and stated “either”. At the end of each trial with a reported change, subjects then reported the direction of perceived change.

### Task Performance

All the subjects learned the change detection task quickly and their performance plateaued after a pre-experimental practice session. Subject performance depended on the informed and uninformed task conditions (**Figure 2**). Similar to results from related experiments^11,12,38^, the hit rate of the informed trials for increasing stimuli was lowest at the smallest stimulus delta (10 Hz), and increased until peaking at the largest delta (50 Hz), with a psychometric threshold (50% hit rate) at a delta of 30.4 Hz (CI: 29.2 to 31.5). This psychometric function shows that subjects were attending to the stimuli. Performance on informed trials followed a similar trajectory for the decreasing stimuli. Hit rates were the lowest at -10 Hz and the highest at -50 Hz. The psychometric threshold for these trials was -27.7 Hz (CI: -28.6 to -26.9).

**Figure 2.**
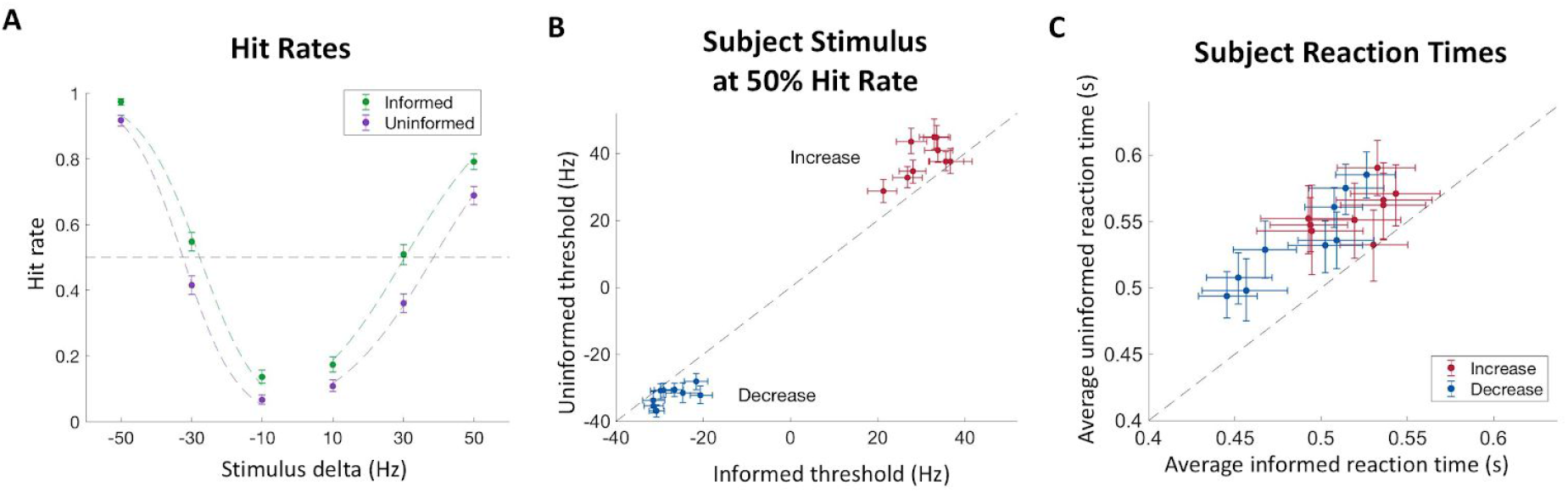
Hit rates and reaction times. A) Performanceplotted as afunction of the change in click rate for each trial condition. Colored dashed lines indicate fits by logistic curves and the dashed line indicates a 50% hit rate psychometric threshold. Data is combined from all subjects. B) The psychometric threshold for each subject in the informed and uninformed conditions of both increasing and decreasing stimuli trials. C) Average hit rate reaction times of each subject for increasing and decreasing stimuli in the informed versus uninformed conditions. All error bars indicate 95% confidence intervals. N=9 subjects.

Performance was noticeably different in the uninformed condition. While hit rates followed the same pattern of increasing with the absolute magnitude of the stimulus change, hit rates were lower than the informed condition for all stimuli, on average. Uninformed trials with increasing stimuli had a psychometric threshold of 38.6 Hz (CI: 37.4 to 39.7), which was significantly higher than informed trials (p < 0.01). Uninformed trials with decreasing stimuli had a psychometric threshold of -32.4 Hz (CI:-33.1 to -31.7), which was significantly larger in magnitude than informed trials (p < 0.01). These differences were consistent across subjects for both the increasing and decreasing stimuli (**Figure 2B**, 7 of 9 subjects and 6 of 9 subjects p < 0.05, respectively).

We also measured the reaction times of hit trials. Consistently, reaction times were longer in the uninformed condition than the informed condition, with subjects responding more quickly to changes when the cue was informative. The average reaction time on uninformed trials was 0.54 s (CI: 0.54 to 0.55), and on informed trials, it was 0.50 s (CI: 0.49 to 0.51), with consistent differences of this type across subjects regardless of whether it was detection of increasing or decreasing stimuli (**Figure 2C**).

Finally, we examined the false alarm rate for each trial type. The average rate was 18.9% (CI: 18.91 to 18.94) and 13.3% (CI: 13.31 to 13.33) for the informed condition, and 11.2% (CI: 11.18 to 11.20) and 5.7% (CI: 5.70 to 5.72) for the uninformed condition, for increasing and decreasing stimulus trials respectively (**Figure 3A**). When plotting the difference between informed and uninformed false alarm rates in **Figure 3B**, we found that in both increasing and decreasing trial groups, the difference was positive (μ: 0.077, CI: 0.063 to 0.092 and μ: 0.076, CI: 0.066 to 0.087, respectively). In other words, the false alarm rates were higher in the informed condition than the uninformed condition. These differences were consistent across subjects for both the increasing and decreasing stimuli (**Figure 3C**, 8 of 9 subjects and 9 of 9 subjects p < 0.05, respectively).

**Figure 3.**
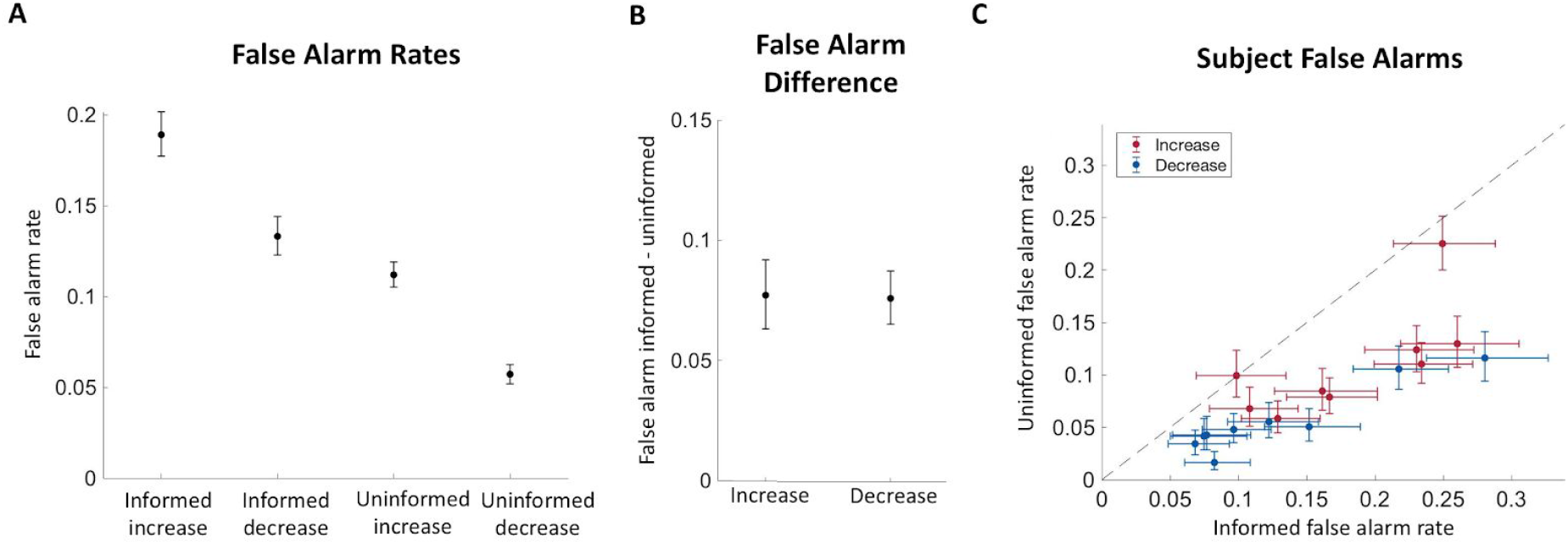
False alarm rates. A) False alarm rate for each trial condition. Data is combined from all subjects. B) Difference between informed and uninformed false alarm rates for combined subject data. C) Informed and uninformed false alarm rates for each subject. All error bars indicate 95% confidence intervals. N=9 subjects.

### Model-Free Analysis

To further assess the differences between trial conditions, we computed the psychophysical reverse correlation (RC) kernels for false alarms of each trial type. The RC kernels were calculated by convolving the click times with a causal half-Gaussian filter and aligning the result to the time of the response for each false alarm trial. This allowed us to reconstruct the average stimulus that preceded a false alarm in order to discern the influence of the stimulus on the false alarm decision. Segments where the RC kernel varies from the baseline click rate indicate the average periods and stimulus magnitudes that influenced the subsequent choice. In our data, increasing false alarms were preceded by an increase in click rate and decreasing false alarms were preceded by a decrease in click rate (**Figure 4A**). The RC kernels also suggest there was a larger change in the stimulus to cause a false alarm in uninformed conditions than in informed conditions, for both increases and decreases. To confirm this difference across subjects, we compared average click rate deviations from the baseline rate over one second preceding a false alarm for each condition (**Figure 4B**). For increasing false alarms, uninformed trials had a larger increase from the baseline stimulus than informed trials (5 of 9 subjects p < 0.05). Similarly for decreasing false alarms, uninformed trials had a larger decrease from the baseline than informed trials (7 of 9 subjects p < 0.05).

**Figure 4.**
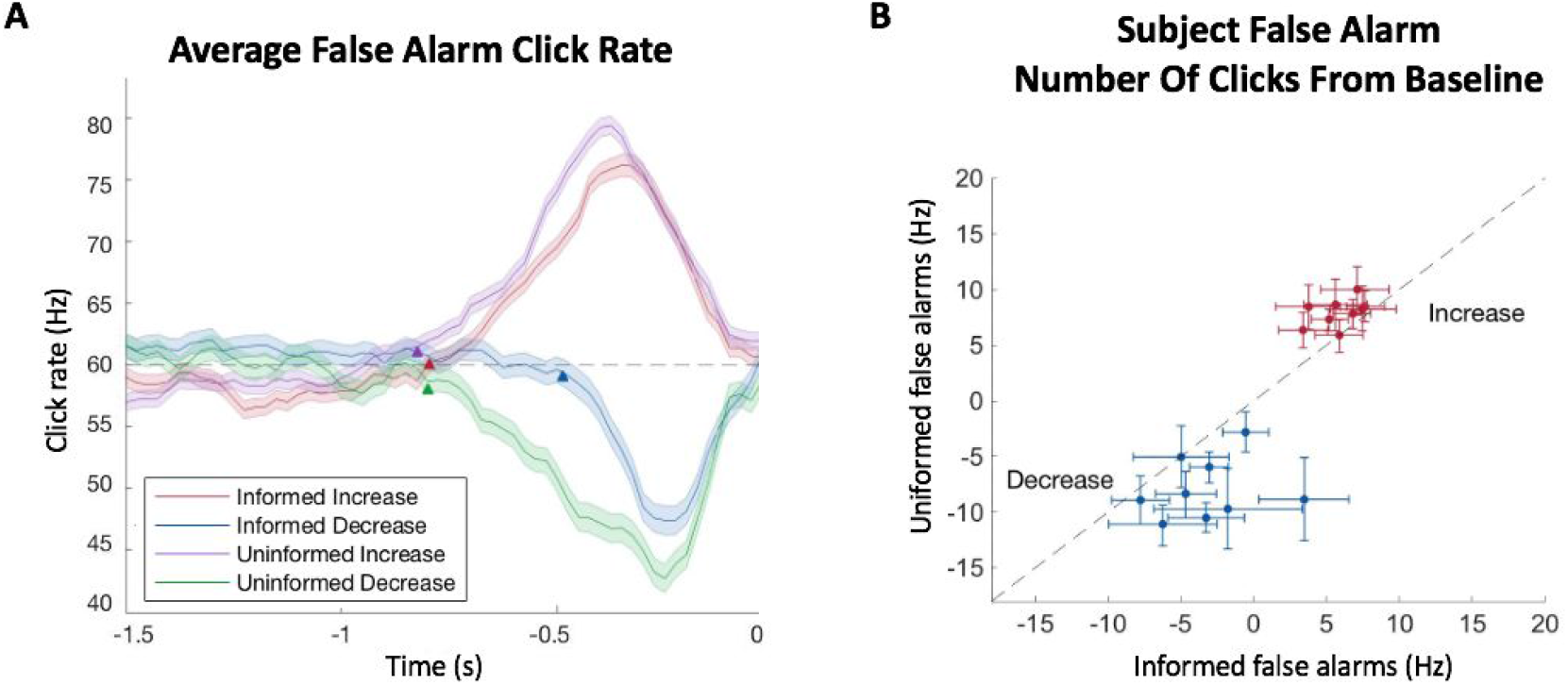
False alarm RC kernels. A) Detection kernels of different trial types for pooled subject data. Shaded regions indicate the standard error of the mean. Arrows indicate the kernel start point. Dashed line indicates baseline click rate. Data is combined from all subjects. B) The average deviation in click rate from the baseline preceding a false alarm for increasing and decreasing trials. Error bars indicate 95% confidence intervals. N=9 subjects.

The starting points of the RC kernels serve as an estimation for the timescale of evidence evaluation. For example, an earlier starting point indicates a longer timescale. We found that the starting points of the RC kernels for increasing trials was slightly shorter in the informed condition at -0.80 s (CI: -0.81 to -0.77) than the uninformed condition at -0.82s (CI: -0.85 to -0.79). In decreasing trials, the start time of the informed condition was significantly shorter at -0.49s (CI: -0.50 to -0.47, p < 0.05) than the uninformed condition at -0.80s (CI: -0.83 to -0.77). In sum, the RC kernels suggested that subjects may adjust both decision bounds and decision filters in the different conditions.

### Model-Based Analysis

To further characterize the behavioral adjustments suggested by our model-free analyses, we fit subject responses to a sequential sampling behavioral model that parametrizes both decision filters and bounds. In general, these models predict a response time for a given stimulus by estimating the value of a decision variable over time and responding when the decision variable crosses a detection bound. Specifically, our model estimates the decision variable by convolving stimulus clicks with an exponential detection filter, and computes the response time by adding a non-decision time component to the bound crossing time to account for sensory and motor delays in responding. Thus our model includes five free parameters: filter width, noise amplitude, bound, non-decision time, and non-decision time variability (see methods). In total, three separate models were fit to trials grouped by cue (informed increase, informed decrease, and uninformed) from both individual and combined subject data. Informed models contained a single process to detect either increasing or decreasing stimuli depending on the cue (**Figure 5A**), whereas uninformed models contained two separate processes to detect both increasing or decreasing stimuli (**Figure 5B**). Each model was fit to the behavioral data by maximizing the likelihood that the subjects’ response times could be produced by the model parameters.

**Figure 5.**
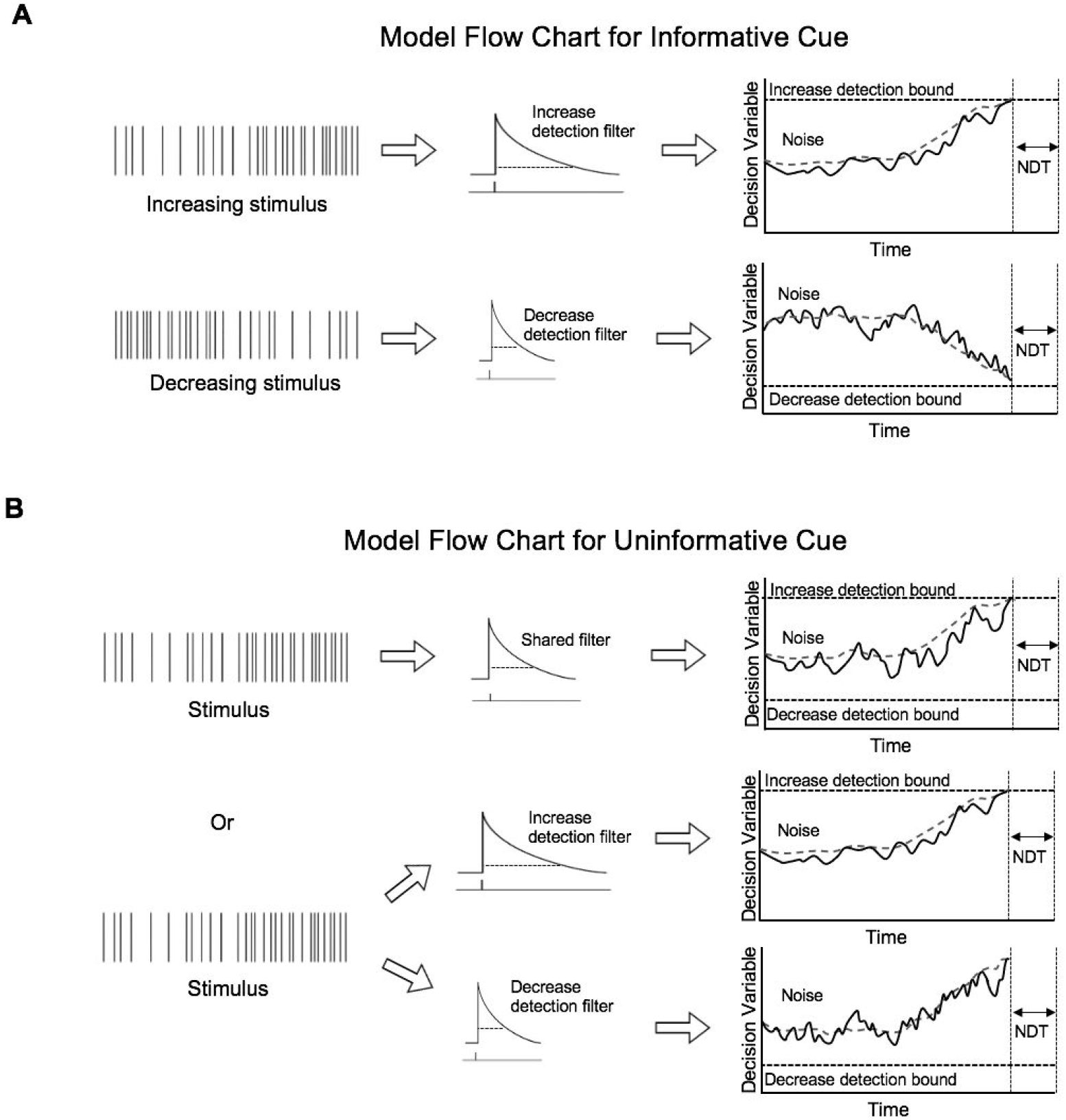
Model schematic. A) Model schematic for informative trials. A decision filter is convolved on a stimulus and decision variable is estimated. When the decision variable crosses the detection bound, a choice is made. B) Model schematic for uninformative trials. The stimulus may be either increasing or decreasing. Either a shared filter can be used regardless of increasing or decreasing stimuli (top) or two different filter widths can be used simultaneously (bottom).

We used this modeling framework to better understand how subjects alter their decision making process when they are informed versus uninformed about the direction of stimulus change. To this end, we asked whether behavior in uninformed conditions was better explained by a model with separate or shared detection filters (**Figure 5B**). To best characterize the difference between these two model configurations, we used two fully-separate single-bound models with each detection filter condition: shared and unshared. By comparing the two conditions in this way, we can isolate the effect of constraining the filter widths because any difference between the two filter conditions cannot be explained by the other model parameters.

Model fit improvement was characterized by comparing log model evidence lower bound (ELBO) values between each fit. ELBO values are analogous to Bayes factors and naturally penalize overfitting, which allows us to compare each model even though they contain different numbers of parameters. The ELBO metric favored the model with unshared filter width parameters for both the combined behavioral data (ELBO difference = 341) and for all nine subjects (mean ELBO difference = 41; Range: 5 to 110). Similar results were found using

Bayesian Information Criterion and Akaike Information Criteria for model comparison. In addition, the shared model produced psychometrics that matched the experimental data very well without explicitly fitting the model to these metrics (**Figure 6A-B**). This suggests that subjects used separate decision making processes that operated simultaneously with distinct timescales of evidence evaluation. Indeed, fits on individual subject data consistently revealed shorter filter widths for detecting decrease changes than those for detecting increase changes (**Figure 6C**).

**Figure 6.**
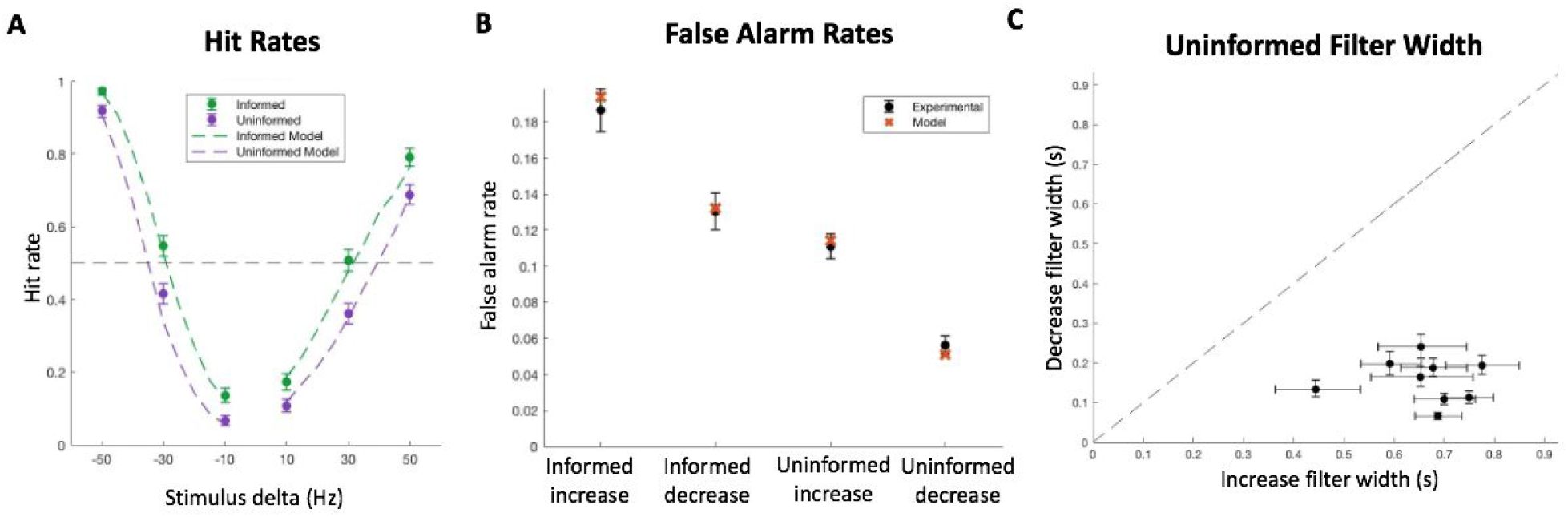
Model Fit. A) Experimental combined subject data with performance plotted as a function of change in click rates compared to the estimated model performance. B) Experimental false alarm rates for combined subject data in each trial condition compared to estimated model false alarm rates. C) Unshared filter width parameter pairs in uninformed models fit to individual subject data. All error bars indicate 95% confidence intervals. N=9 subjects.

To better understand how subjects adjusted their decision process when cued to the nature of the change, we compared how the four major model parameters (filter width, bound, noise amplitude and non-decision time) varied between informed and uninformed conditions for each change detection process (increase or decrease). For increasing stimuli, we found that the primary change in the decision process could be described as a change in the detection bound. The model fit to combined subject data showed a significant decrease in the bound when subjects were informed (83.9 Hz versus 78.3 Hz, p < 0.01) (**Figure 7A**), which is consistent with our model-free analysis. Additionally, models fit to individual subject data revealed the same trend in eight of the nine subjects (**Figure 7B**). Although other parameters seemed to vary slightly between conditions when fit to the combined subject data (such as filter width, **Figure 7A**), these differences were not as consistent across subjects (**Figure 7C**). For decreasing stimuli, we found that the change in the decision process was more complex, as the filter width, noise, and bound all seemed to change by a substantial amount (**Figure 7D**). In contrast to the increasing stimuli, the change in the filter width was consistent across subjects for decreasing stimuli, but the change in bound was not (**Figure 7E-F**). Further comparing differences between increase and decrease model parameters, there seems to be a robustly different set of optimal values depending on the change direction, which can be most strongly seen in the filter width being substantially longer for increasing stimuli. This suggests that optimizing the ability to detect changes in different directions requires different parameter tunings. With this understanding, we found that uninformed decrease change detection parameters were generally pulled away from their optimal informed values toward their optimal increase detection values, which can be seen especially strongly by the increased filter width (compare **Figures 7D** with **7A**).

**Figure 7.**
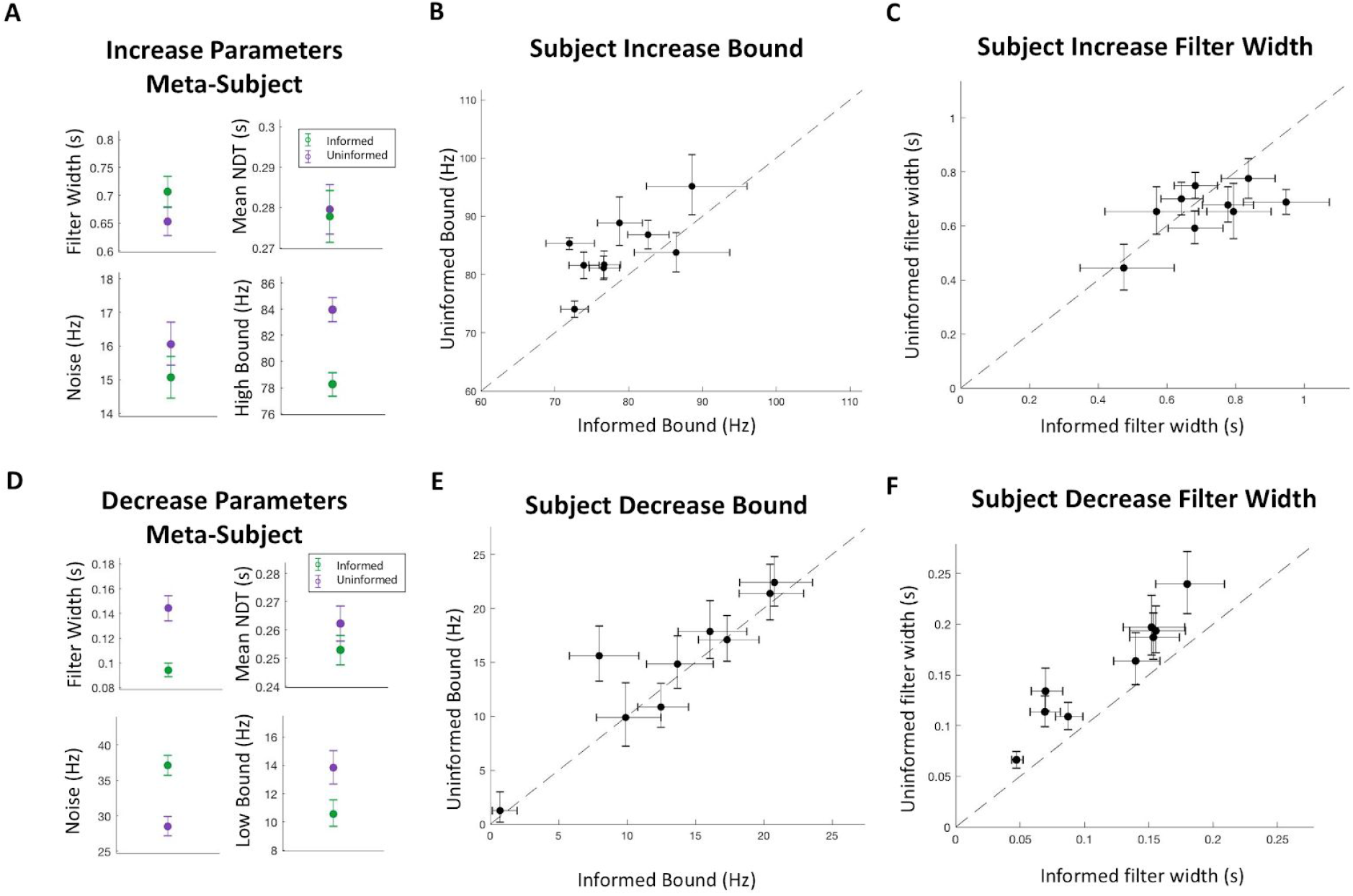
Model Parameters A) Increase change detectionparametersfor the informed and uninformed conditions fit to combined subject data. B) Increase bound values between informed and uninformed conditions from models fit to individual subject data. C) Increase detection filter widths between informed and uninformed conditions from models fit to individual subject data. D) Decrease change detection parameters for the informed and uninformed conditions fit to combined subject data. E) Decrease bound values between informed and uninformed conditions from models fit to individual subject data. F) Decrease detection filter widths between informed and uninformed conditions from models fit to individual subject data. All error bars indicate 95% confidence intervals. N=9 subjects.

## Discussion

We used a stochastic auditory change point detection task to investigate how knowledge about the space of possible changes affects human change point detection. In our task, changes to the generative click rate in either direction occurred at unpredictable times, and subjects received either an uninformative or an informative cue about the direction of the impending change. Through model-free and model-based analysis we found two primary conclusions: subjects use knowledge about the dimension of the change to narrow their ‘decision criteria’, and subjects are able to process the same sensory information over multiple distinct timescales.

From the behavioral results, subjects were able to discern changes at a greater rate and in a shorter amount of time when they were informed of the direction of the change (**Figures 2A,C**). However, this improved performance came at the cost of an increase in false alarms (**Figure 3**). When comparing the false alarm trials between conditions with psychophysical reverse correlation analyses, we found that subjects responded to smaller deviations from the baseline click rate when they were informed about the direction of the change (**Figure 4**). This suggests that subjects narrowed their ‘decision criteria’ when they were able to reduce the number of attended dimensions in the stimulus stream. By reduced ‘decision criteria’, we imply that subjects required less evidence to commit to their decision, which meant subjects were able to react faster and were more likely to respond to changes when they did happen. Further supporting this claim, a reduced evidence threshold would make subjects more likely to falsely respond to stimulus noise, which we also found to be the case. The question still remained, however, how this reduced bound can be implemented. Intuitively, we can imagine three main non-mutually exclusive implementations that could account for this behavior: improving the ability to remember past stimuli to compare with current stimuli to better discern changes, reducing the amount of evidence required to suggest that the new stimuli are different from past stimuli, and increasing the ability to filter out stimulus noise while making this comparison. To better understand how these implementations might explain our behavioral results, we developed a model that incorporates terms for all of these scenarios (filter width, bound, and noise, respectively) and fit it to our data.

When comparing best-fit model parameters between informed and uninformed conditions, we generally found what looked to be different implementations depending on the direction of the change. One consistency between directions is that our models suggest subjects were not able to improve the signal-to-noise ratio of the underlying evidence, as characterized by our noise term. Rather than being able to directly increase perceptual sensitivity, subjects seemed to improve their change detection capacity with a mixture of adjustments to their decision bounds and timescales of evidence evaluation. This is consistent with other studies suggesting an important role of changes in the decision criterion for improvements in performance that occur with attentional cues^28,29^. This is important because improvements in accuracy with cuing are often ascribed to attentional processes sometimes thought to improve perceptual sensitivity^20,21^. We do not rule out that improved signal to noise can be important for change detection behavior, but it is interesting that we find no evidence for it, even with the cuing manipulation that we perform. An implementation that was unique for cues informing increases was that subjects reduced their decision bound, consistent with the inferences drawn from our model-free analyses. This suggests that subjects reduce the amount of evidence required to commit to the decision when there are fewer alternatives to detect and is reminiscent of the reductions in decision bounds found in discrimination tasks with fewer alternatives to consider^39^. Supporting this claim, neural correlates that are potentially consistent with higher decision thresholds when considering multiple alternatives^40^ have been reported in parietal cortex^39,41,42^, frontal cortex^43,44^, and subcortical regions involved in action selection^45,46^. However, none of those studies involved change point detection, so it would be interesting to explore whether similar neural mechanisms may be involved in our task. For decreasing changes, the model suggested that all three decision criteria parameters were adjusted when subjects were informed. From our results, we are not able to conclude that any one change was more important than the other, but in general each of the parameters moved in a direction toward their best fit values for cued increase trials when subjects were uninformed. This result seems to suggest that there may be some resource sharing between the two processes when both change directions were attended to simultaneously.

When comparing parameters between informed conditions, we found that there were distinct differences in the best fit values, especially in regard to filter width. We speculate that this difference may relate to an important asymmetry for detecting increase changes versus decrease changes in our task, as can be seen in the behavioral psychometrics. In terms of our model, the decision variable moves closer to the increase decision bound in a stepwise manner with each click, and may therefore be more susceptible to noise and explain higher false alarm rates for detecting increases. In contrast, the decision variable smoothly decays toward the decrease decision bound in the absence of clicks, and may therefore be more robust to noise and explain lower false alarm rates for detecting increases. This asymmetry can best be seen in the model’s filter width terms. In order to respond quickly to decreases, it makes sense that the decision value should decay toward the decrease bound faster than when detecting increases, which we found to be the case in our model fits. Additionally, a longer filter width smooths the decision value’s upward trajectory thereby moderating the influence of stimulus noise, such as short bursts of clicks. With this understanding of the differences between increase and decrease detection processes, we investigated whether the data from the uninformed condition could be best described by two detection processes with a shared filter width, or by two completely separate detection processes, each with their own timescale of evidence evaluation. We found that the latter best explained our data, which suggests subjects were able to evaluate the same stimuli over multiple timescales simultaneously.

This finding closely mirrors results from a previous experiment in our lab where confidence in a change point detection had a longer timescale of evidence evaluation than the timescale required to detect the change^12^. Additionally, our finding that subjects can flexibly adjust their timescales of evidence evaluation on the same stimuli is supported by previous work from our labs and others. For example, in the visual and auditory domains, humans adjust their timescale of evidence evaluation to match the distribution of signal durations and timing they experience^10,38,47–49^. Humans can also adjust their timescales of evidence evaluation when classifying the location of visual stimuli, using a shorter timescale in more volatile environments and a longer timescale in more stable environments^9^. Likewise, both humans and rats can adjust their timescales of evidence evaluation in a similar manner when discriminating the current state of a changing sensory stimulus, using a shorter timescale when the stimulus changes more often and a longer timescale when it changes less often^8,9^. A distinct difference between the prior studies and ours is that all of these previous findings derived from tasks that involved a timescale of evidence evaluation that varied over blocks of trials, giving subjects time to adapt to a new optimal evaluation period. In contrast, we demonstrate that subjects were able to rapidly change their timescales on a trial-by-trial basis, and additionally, were able to simultaneously process the same sensory stream with multiple timescales when needed.

The capacity to simultaneously evaluate evidence along multiple timescales provides insight into potential underlying neural mechanisms. It seems to rule out mechanisms that allow only a single timescale of evaluation for a particular source or stream of evidence, even if that timescale is tunable. Instead, any proposed neural mechanism must be able to support at least two distinct timescales. One potential mechanism for this is to have distinct neural populations with separately tunable timescales that process the same stream of evidence in parallel. Another mechanism is a single population of neurons that contains a heterogeneity of timescales^50–54^, which is suggested by a number of network architectures^55,56^. One advantage of these architectures is that they readily support evidence evaluation across multiple timescales without information loss or re-tuning of the networks involved in the evaluation process.

Overall, our data supports the idea that parallel evaluation of the same stream of evidence can occur along distinct timescales for change detection with multiple alternatives. Furthermore, cued attention narrows the decision space and allows observers to make changes in decision processes that improve performance. These results establish three important capacities of information processing for decision making that any proposed neural mechanism of evidence evaluation must be able to support: the ability to simultaneously employ multiple timescales of evidence evaluation, the ability to rapidly adjust those timescales, and the ability to modify the amount of information required to make a decision in the context of flexible timescales.

## Author Contributions

All authors were involved in conception and design of the experiments. AB, TS, CM, and MR were involved with data collection. AB and TS led the data analysis with input from TDH. AB drafted the manuscript with assistance from TS and TDH. All authors reviewed the final manuscript.

## Acknowledgements

We thank Preetham Ganupuru and Adam Goldring for helpful feedback and comments throughout all phases of this study.

## Notes

### Competing Interest Statement

The authors have declared no competing interest.

